# *Anopheles azevedoi* (Ribeiro, 1969) in Angola: New geographical records, molecular characterization, and insecticide susceptibility profile of a commonly misidentified species

**DOI:** 10.64898/2026.02.22.706856

**Authors:** Gonçalo Alves, Catia Marques, Paula Marcet, Vicente Chipepa, Anya Fedorova, Alice Sutcliffe, Joana do Rosario, Dinorah Calles, Arlete Dina Troco, Moisés Sacamário Chissanga, Félix Espalhado, Teresa Nobrega, Carla Sousa, João Pinto, Cani Pedro Jorge, José Franco Martins, Melissa Yoshimizu, Carolina Torres Gutierrez

**Author notes:** Corresponding authors: Gonçalo Alves and Carolina Torres Gutierrez. These two authors equally contributed as first authors (Gonçalo Alves, Catia Marques).

## Abstract

**BACKGROUND:** Angola ranks among the five countries with the highest malaria burden globally. The Ministry of Health in Angola has consistently partnered with international donors to sustain entomological surveillance and vector control strategies in a context of high malaria burden.

**METHODS:** Vector surveillance was carried out in Luanda, Benguela, Namibe and Cuanza Sul provinces from 2016-2022. Collected adult mosquitoes were tested to assess the presence of *Plasmodium* parasites and determine blood sources. Larvae collections provided live material to test insecticide susceptibility in local *Anopheles* populations. Taxonomic determination of mosquitoes was based on external morphology and confirmed with molecular assays. The presence of *Anopheles azevedoi* was confirmed through morphology and genetic sequences, and errors in the original species determination were detected, discussed and corrected.

**OBJECTIVES:** The study aimed to update the geographical range of *Anopheles azevedoi* in Angola and monitor the species’ susceptibility to public health insecticides.

**FINDINGS and MAIN CONCLUSIONS:** We report on populations of *Anopheles azevedoi* occurring along the western coast of Angola, a highly abundant species with anthropophilic behavior in urban areas. *Anopheles azevedoi* is widely resistant to pyrethroids and DDT but fully susceptible to chlorfenapyr. We contribute with *COI* and ITS-2 barcoding sequences for future species identification and explain the reasons for which this species has been for long misidentified in Angola.

## INTRODUCTION

Angola is in the western coast of Southern Africa and is a young country that gained independence in 1975, after being a Portuguese colony for hundreds of years (1). Despite universal and free health care, most of the Angolan population lives in households with limited resources as it is described by the wealth index in demographic and health surveys (2, 3).

According to the Malaria Indicator Survey the “Angolan population is fairly young as a result of a demographic regime of high fertility and high mortality” (4). Life expectancy in the country is among the lowest in the world, with sharp socioeconomic inequalities and high infant mortality rates; with 16 neonatal deaths out of 1,000 live births, and infant mortality documented at 32 deaths per 1,000 live births (2, 5). Angola is among the five countries with the highest malaria burden, accounting for 3.4% of global cases and 3.2% of global deaths (6). Malaria represents the leading cause of death in Angola and is responsible for high maternal and infant morbidity; it is also the main illness diagnosed after medical consultations, and the primary reason for work and school absenteeism (3, 4). In this context, the Ministry of Health of Angola (MoHA) has sustained long and strong partnerships to support malaria surveillance, prevention, and control in Angola since 2004. The Angolan National Malaria Control Program (NMCP), in partnership with Global Fund, MENTOR, and the President’s Malaria Initiative (PMI), has operated in different provinces with the highest malaria incidence and led activities associated with vector control, case management, supply chain of public health commodities, prevention of malaria in pregnancy, social and behavior change, surveillance, monitoring, and evaluation.

Vector surveillance programs over the past twenty years in Angola have confirmed the presence of several malaria vectors included in the *Anopheles gambiae* complex and the *An. Funestus* group (7) such as *An. coluzzii*, *An. arabiensis*, *An. melas*, and *An. funestus* s.s. (8–14). *Anopheles azevedoi* was first reported in Angola in 1969 by Henrique Ribeiro in the city of Moçâmedes, located in Namibe Province (15). Following this initial report, the species was subsequently recorded in several locations along the southern coastal regions of the country (14, 16). Within the current classification of the genus *Anopheles*, *An. azevedoi* belongs to the subgenus *Cellia*, series Paramyzomyia, group Cinereus (17). According to Ribeiro (16), this Angolan species is closely related to *An. listeri* and *An. multicolor* with great morphological and ecological similarity. A few species within Paramyzomyia, such as *An. azevedoi*, An. *listeri* and *An. multicolor*, are recognized in the literature by their tolerance to salt-water during their immature phases, which occur in dry biomes, with documented behavior as anthropophilic and exophilic populations (16). This study provides new records of *An. azevedoi* distribution in the country, its insecticide resistance profile and reports, for the first time, its molecular sequencing identification.

## MATERIALS AND METHODS

### Study Area

Angola is located on the Western coast of Southern Africa sharing borders with the Republic of Congo and the Democratic Republic of the Congo in the North, Zambia to the East and Namibia in the South. Angola’s climate is characterized by hot humid summers and mild dry winters (18). Vast areas of the country are covered by savanna, shrubland, grassland, tropical forest or bare land. Most of the cropland regions are located in Angola’s central plateau, with elevations above 1000 m.

The country is divided into 21 provinces and its capital is Luanda, the largest city in the country, with an estimated population of over 8 million, with densely populated urban slums (2). Luanda is a maritime port of great importance to the Angolan economy. The main economic activities include oil refinery and diverse manufacturing and agricultural processes such as beverages, textiles, cement, metallurgy and car assembly (1).

Benguela, a province located in the west region of Angola, lies along the Atlantic coast. This province occupies a total area of 39,508 km^2^ with altitudes ranging from sea level to 2,547 m towards the east (19). It is a steppe coastal plain dominated by a vegetation of shrubs and herbaceous taxa. Benguela’s climate is tropical, and it is conditioned by strong dry offshore winds (20).

Cuanza Sul is in the central-west region of Angola with a total area of 55,660 km^2^. The western section of the province lies along the Atlantic Ocean. This Angolan region is characterized by a diverse geography encompassing river valleys, hills of varied altitude, and coastal plains (20).

The province of Namibe is a southern region with vast desertic areas which serve the country as one of the main maritime ports and a place of developed artisanal and industrial fishing (1).

### Mosquito sampling

Sampling sites were selected by the National Malaria Control Program (NMCP) based on the documented malaria intensity and history of insecticidal use in different regions of Angola. Mosquito collections were conducted by two independent teams led by the NMCP in collaboration with Populations Services International (PSI)/PMI and MENTOR partners. PSI/PMI conducted activities in the provinces of Luanda, Namibe and Benguela; and MENTOR was based in Benguela and Cuanza Sul. Researchers of Institute of Hygiene and Tropical Medicine (IHMT) from Nova University of Lisbon (NOVA), participated as consultants for PSI. Sample locations are detailed in Table I.

**Table I.** Geographical location and characteristics of sampling areas in different provinces of Angola.

| Province/<br>Field team | District | Village or<br>site/year of<br>collection | Geolocation<br>(Latitude,<br>longitude) | Type of breeding site<br>(Permanent or Temporary) | Vegetation | Quality of<br>water | Type of<br>collection/<br>Phase |
| --- | --- | --- | --- | --- | --- | --- | --- |
| Benguela | Catumbela | Multiple sites,<br>2015 | Not recorded | Roadside drains, brick/sand pits,<br>puddles, hoof prints, rice fields | Variable/ details<br>not available | Stagnant | Larvae; Phase-I |
| Namibe | Namible | Multiple sites,<br>2016 | Not recorded | Roadside drains, brick/sand pits,<br>puddles, hoof prints, rice fields | Variable/ details<br>not available | Stagnant | Larvae; Phase-I |
| Luanda | Cacuaco | Paraíso, 2021 | -8.797,<br>13.34505 | Roadside pools (temporary) | Dense grass<br>along the road | Clear | Larvae; Phase-II |
| Luanda | Ingombotas | Ilha de Luanda,<br>2016, 2022 | -8.79172,<br>13.22733 | Abandoned swimming pool<br>(Hotel Panorama - close to<br>coastline, permanent) | Grown trees<br>around<br>swimming pool | Stagnant water<br>and industrial<br>debris. | Larvae and<br>adults; Phase-I<br>and Phase-II |
| Benguela | Benguela | Riacho, 2022 | -12.60492,<br>013.37751 | Natural / riverbank (permanent) | Semi-arid | Polluted | Larvae;<br>Phase-III |
|  |  | Bairro Ndacota,<br>2022 | -12.59659,<br>013.40367 | Puddles (permanent) | Semi-arid | Polluted | Larvae;<br>Phase-III |
|  |  | Rio Cavaco, 2022 | -12.56944,<br>013.41986 | Wells dug in the riverbank<br>(temporary) | Semi-arid | Clear | Larvae;<br>Phase-III |
| Benguela | Catumbela | São Pedro, 2022 | -12.44987,<br>013.53941 | Puddles (permanent) | Grass patch | Polluted | Larvae;<br>Phase-III |
| Cuanza Sul | Sumbe | Feira, 2022 | -11.17840,<br>013.85678 | Puddles (semi-permanent) | Present | Clean | Larvae;<br>Phase-III |
|  |  | Carimba, 2022 | -11.07662,<br>013.85802 | Artificial / Water tank<br>(temporary) | Absent | Turbid, dark | Larvae;<br>Phase-III |

Field labor under entomological surveillance efforts was conducted through monthly community-based vector sampling. Technicians conducted visual search for potential breeding sites around populated zones of the designated areas to collect immature forms (larvae and pupae) to determine species composition and distribution, as well as to conduct insecticide susceptibility testing. Municipalities included as designated areas were selected by the NMCP according to their epidemiological classification, as high malaria transmission risk, and presence of a provincial health office belonging to the Ministry of Health’s network. Larval collections took place during different timeframes: Phase-I sampling took place between 2015 and 2016 in the provinces of Namibe and Benguela, Phase-II sampling took place between 2020 and 2022 in the province of Luanda and Phase-III sampling occurred in the provinces of Benguela and Cuanza Sul from March to May 2022.

Immature *Anopheles* were collected using 350 ml standard mosquito dippers (BioQuip Products, Rancho Dominguez, CA, USA) and transported from larval habitats to the corresponding insectaries closest to each province. Personnel collected as many larvae as possible in each breeding site (variable number of dips depending on size of sites). The space hosting the insectary in Luanda corresponded with the Instituto Nacional de Investigação em Saúde (INIS/ National Institute for Health Research), and an insectary facility owned by NMCP in the province of Benguela was employed by the MENTOR team. For other provinces, field teams used different locations to establish field insectaries where larvae and pupae were reared in their breeding water and newly emerged adults kept in cages under a photoperiod of 12 h light:12 h dark, ambient temperature of 28 ± 2 °C, and a humidity of 75 ± 5%. Reared female specimens were employed for monitoring insecticide susceptibility.

Adult sampling was carried out only in Luanda near Ilha de Luanda (Ingombotas/Luanda). Field teams employed CDC light traps (CDC LT) inside houses from 6 p.m. to 6 a.m. nearby where breeding sites had been previously identified. Verbal consent from house-owners was obtained. Adult mosquitoes were initially sorted and identified and a subsample of such material (n=49) was sent to the Centers for Disease Control and Prevention (CDC), Entomology Branch (Atlanta, GA, USA), for species confirmation through morphological and molecular analyses. Field specimens were identified by using characters of their external morphology following taxonomic keys (21–24).

### Molecular Identification of species, infectivity studies and blood meal analysis

A subsample (n=49) of the adult specimens collected in Luanda were sent to the Entomology Branch facilities at the Centers for Disease Control and Prevention (CDC) in Atlanta, Georgia, United States, for species confirmation through PCR amplification and DNA barcode sequencing.

DNA extraction was carried out using Extracta^TM^ DNA Prep following manufacturer recommendations (Quantabio, Beverly, MA, USA). Molecular species identification of the *Anopheles gambiae* complex and *Anopheles funestus* group followed previously described PCR-based protocols and considerations (25–27).

Specimens that gave an inconclusive result with the *An. gambiae* or *An. funestus* species-specific PCR assays (i.e. no reliable amplification bands) underwent Sanger sequencing in an ABI 3500 Prism Genetic Analyzer (Applied Biosystems) automated sequencer using the ITS-2 and *COI* barcoding genes following procedures described by published studies (28, 29). Sequences were assembled and analyzed using Lasergene Seqman Pro (DNASTAR, Inc.).

Comparison analyses for species identification using genetic sequences were performed using Basic Local Alignment Search Tool (BLAST; National Center for Biotechnology Information, Bethesda, M.D.) (30). The criteria for species identification were DNA sequence similarity, with values of 98% similarity or higher, as indicative of same species. DNA sequences for both genes were blasted against the complete barcoding database available in GenBank [https://www.ncbi.nlm.nih.gov/genbank/]. The sequences were also compared against a dataset of existing *Anopheles* samples from Angola that had been kept in the CDC facilities since the time of Phase-I sampling in 2015 (unpublished observations).

To assess natural infectivity of *Anopheles* specimens, a sample of adult mosquitoes (n=33) captured in Luanda sites as adults by CDC LT were screened for infective *Plasmodium* parasites using a circumsporozoite ELISA for *P. falciparum* using the head and thorax of each specimen (31). The remaining body parts were used for molecular species identification. Moreover, a more sensitive bead-based multiplex immunoassay (32) that allows for simultaneous screening for *P. falciparum* and *P. vivax*, was run in parallel. Blood meal analysis for these specimens was assessed by screening the DNA extracted from adult specimens with engorged abdomens using PCR amplification and sequencing of the mitochondrial cytochrome b marker (33).

### Insecticide susceptibility testing

The insecticides prioritized for the susceptibility testing respond to those active ingredients that have been included in malaria control strategies in Angola, including indoor residual spraying (past strategies) and insecticide-treated bed nets (ITNs) which were recently distributed by the NMCP in different regions of the country.

Insecticide testing was conducted with impregnated papers using WHO tube or bottle bioassays and following WHO guidelines (34, 35). All insecticide susceptibility tests were performed at 25 °C ±2 °C and 70 % ± 10 %. WHO test tubes involved impregnated papers at specific concentrations including the discriminating concentration (or diagnostic dose) (1x) and intensity concentrations (5x and 10x) (34, 35). Tests were performed with 3-5-days old non-blood fed monospecific female mosquitoes, which emerged from immatures collected from wild breeding sites (Table I). A sample of a minimum of 100 mosquitoes were tested per insecticide to assess susceptibility to the discriminating concentrations of deltamethrin (0.05%), permethrin (0.75%), alpha-cypermethrin (0.05%) and fenitrothion (1.00%). In addition, pyrethroids were tested in combination with the synergist piperonyl butoxide (PBO), according to the WHO guidelines (34–36). For the synergist assays mosquitoes were pre-exposed to PBO (4%) for one hour before being exposed to the pyrethroid insecticides for another hour. Two tubes lined with silicone oil treated papers were set in parallel during each test and served as negative controls with at least 20 females per control tubes. All exposures were done for one hour after which mosquitoes were transferred to resting tubes. Mortality was determined 24 hours after exposure as the number of dead mosquitoes divided by the total tested for each insecticide (35, 36).

To evaluate the susceptibility to chlorfenapyr, WHO bottles were coated with the diagnostic concentration (100 ug/bottle) following the standard protocol (37). Mortality was recorded 24, 48, and 72 hours after exposure. Two bottles coated with acetone were prepared similarly to serve as negative controls. Tested mosquitoes were identified to species or species group by morphological characters using taxonomic keys (21–24).

### Data analyses and geographical records

Mortality values after exposure to different insecticides were recorded and graphed in Excel (Microsoft Windows 10); and maps depicting the known distribution of *An. azevedoi* were generated with QGIS geographic information system (http://www.qgis.org/).

### Taxonomic consultation

A small number of mosquitoes (n=10; Phase-I and Phase II) was sent to the previously known Walter Reed Biosystematics Unit (WRBU/ Smithsonian Institution, National Museum of Natural History, Washington D.C) requesting a taxonomic confirmation of female and male specimens.

## RESULTS

An overview of geographical origin and findings from our collaborative work has been summarized in Table II. The bulk of the biological specimens were employed for the testing of insecticide susceptibility in the different provinces where entomological surveillance was led by Angola’s NMCP.

**Table II.** Description of specimens collected from Angola and procedures performed with the biological material. (*) biological material obtained during Phase-I sampling (PSI/PMI) in Angola; transported and stored in the CDC bio bank and processed in 2021 when additional comparisons with Angolan samples were needed for confirmation of molecular results of 2021-2022 data. This was done to confirm the presence of *Anopheles azevedoi* in Angola from material coming from earlier independent collections.

| Geographic origin (Province) | Total number of specimens | Result | Original taxonomic ID vs. Corrected ID. |
| --- | --- | --- | --- |
| Benguela | Larvae reared to adults*; n=209 | DNA sequences (*COI*, *ITS2*). | Originally labeled as *An. gambiae* s.l. 169 specimens corresponded to *An. azevedoi*. Additional taxonomic identifications not included in this study. |
| Namibe | Larvae reared to adults*; n=110 | DNA sequences (*COI*, *ITS2*). Taxonomic confirmation by Smithsonian (specimens donated as vouchers). Sequences submitted to GenBank. | Originally labeled as *An. gambiae* s.l. 38 specimens corresponded to *An. azevedoi*. Additional taxonomic identifications not included in this study. |
| Luanda | Larvae reared to adults /n=49 | DNA sequences (*COI*, *ITS2*); monitoring insecticide susceptibility. | Originally labeled as *An. parensis*; 48 specimens were identified as *An. azevedoi*, 1 specimen as inconclusive. |
| Luanda | Adults/n=33 | *Plasmodium* species were not found in screened adults. 3 adult specimens were found with human blood. | 26 specimens confirmed as *An. azevedoi*. Additional taxonomic identifications (other species), not included in this study. |
| Luanda | Larvae reared to adults/n=1,521 | Morphological ID; insecticide susceptibility profile for pyrethroids and pyrrole. | Originally identified as *An. azevedoi*/*An. listeri*. Corrected as *An. azevedoi*. |
| Benguela | Larvae reared to adults/n=890 | Morphological ID; Insecticide susceptibility profile for pyrethroids and DDT. | Originally identified as *An. azevedoi*/*An. listeri*. Corrected as *An. azevedoi*. |
| Cuanza Sul | Larvae reared to adults/n=178 | Morphological ID; Insecticide susceptibility data for pyrethroids. | Originally identified as *An. azevedoi*/*An. listeri*. Corrected as *An. azevedoi*. |

### Taxonomic identity of specimens

Records of the morphological identification of Angolan specimens from four different provinces varied according to the taxonomic knowledge of the field technicians. A portion of the specimens were originally misidentified as *Anopheles gambiae* s.l., *Anopheles parensis* or simply as unknown. The most accurate morphological identifications of specimens reached the dichotomic ending of *An. azevedoi*/*An. listeri*, as included in the corresponding taxonomic keys (24). The steps leading to this determination in the most recent key (24) include the following characters: 1) pale and dark areas on wings about equally distributed, base of costa dark or interrupted by scattered pale scales and 2^nd^ main dark area of wing vein 1 with one pale interruption, and 2) maxillary palpus with three pale bands, a basal pale band of maxillary palpus that is much shorter than median band, and dark tip of palpus.

Molecular identification of mosquitoes from Phase-I collections from Angola generated genomic sequences for two genes, *COI* and ITS2. The newly generated sequences were made available on Genbank under the accession numbers OP225396 (ITS2 marker; 633 bp) and OP223465 (*COI* gene; 711 bp). The sequences were evidence that the original identifications of specimens indicating *An. gambiae* s.l. and *An. parensis* were mistaken. After analyzing both morphological and molecular evidence, CDC researchers reached a likely determination of specimens as *An. azevedoi*. The latest known report of *An. azevedoi* in Angola had been by Henrique Ribeiro (16).

The next logical step was to pursuit a consultation with the Walter Reed Biosystematics Unit (WRBU/Smithsonian Institution) submitting female and male specimens of the likely samples of *An. azevedoi*. Morphological studies by the WRBU confirmed that the query specimens were *An. azevedoi*. The availability of male individuals allowed the entomologists at the museum to distinguish between two possible species previously described in Angola, *An. azevedoi* and *An. listeri*, of which females are difficult to distinguish morphologically (21, 23). Reference vouchers for both sexes were deposited at Smithsonian’s facilities in 2022. The information regarding the presence of *An. azevedoi* in the western provinces of Angola was shared with all public health authorities at the national and provincial level within the NMCP during technical meetings held between 2021 and 2022. In addition, CDC researchers delivered trainings to strengthen morphological determination by field personnel. Later in time, when specimens from Phase-II collections (from Luanda) reached the CDC for molecular confirmation, researchers were already aware of the finding of *An. azevedoi* and had DNA sequences to corroborate the presence of this species in the samples. CDC researchers analyzed all steps in the taxonomic keys based on morphological characters and recommended additional guidelines for the field team in Angola.

### New geographical records of *Anopheles azevedoi*

We are updating the known distribution of *An. azevedoi* in Angola (Figure 1). Our results add to the previously documented distribution of the species (15, 16) and show a consistent range along the western coast of the country, with records in the inland territory of Benguela (14). It is worth noting that three sites distributed in Luanda (Cacuaco and Ingombotas) and Benguela (Catumbela) correspond to new geographical records of *An. azevedoi* in Angola. This fact underscores the taxonomic gap observed in previous surveillance efforts, between 1974 and 2024, when this species was consistently misidentified by field teams.

**Figure 1.**
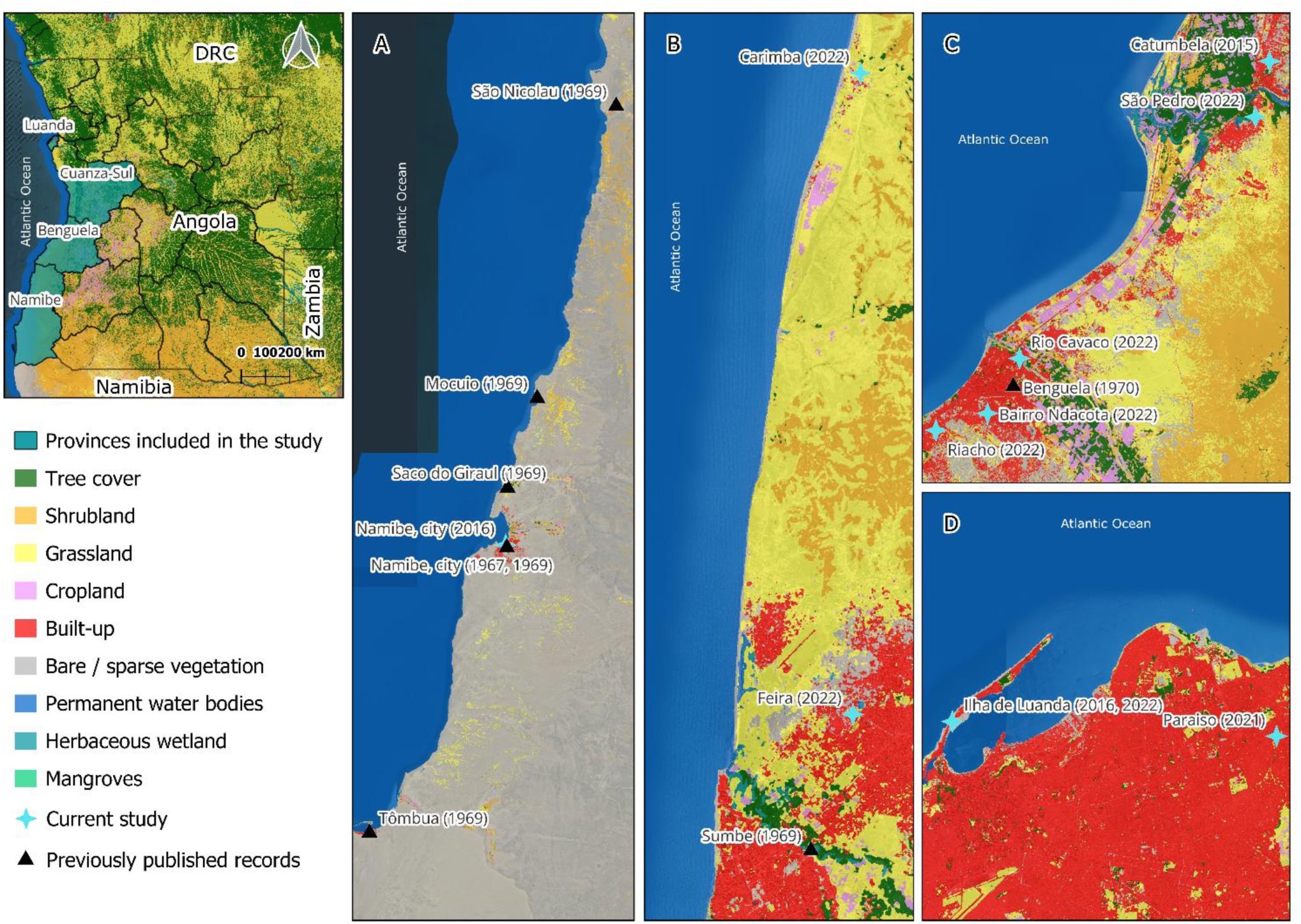
Updated distribution of *Anopheles azevedoi* in Angola. Geographical records from this study are marked with a black triangle (provinces of Luanda, Benguela, Namibe and Cuanza Sul); and other collections in published works (14–16) are presented with a light blue star (provinces of Namibe and Benguela).

### Insecticide susceptibility profile

*An. azevedoi* populations from all study sites across surveillance timelines (2016, 2021, 2022), consistently showed a pattern of resistance to the main pyrethroids and organophosphate insecticides tested, i.e. permethrin, deltamethrin, alpha-cypermethrin and fenitrothion (Figures 3-6).

For the *An. azevedoi* populations from the province of Luanda exposed to alpha-cypermethrin and deltamethrin, with pre-exposure to piperonyl butoxide (a synergist also referred to as PBO), resulted in the restoration of full susceptibility of the tested individuals to these pyrethroids (Figures 3 and 4). For deltamethrin, the restored susceptibility values (99% and 100% mortality) were recorded in mosquito populations from two separate villages during two consecutive years of testing, 2021 and 2022. Similarly, populations exposed to the discriminating concentration for alpha-cypermethrin (0.05%) showed that only the population from Cacuaco village in 2021 recovered full susceptibility to this pyrethroid when pre-exposed to PBO (Figures 2 and 3).

**Figure 2.**
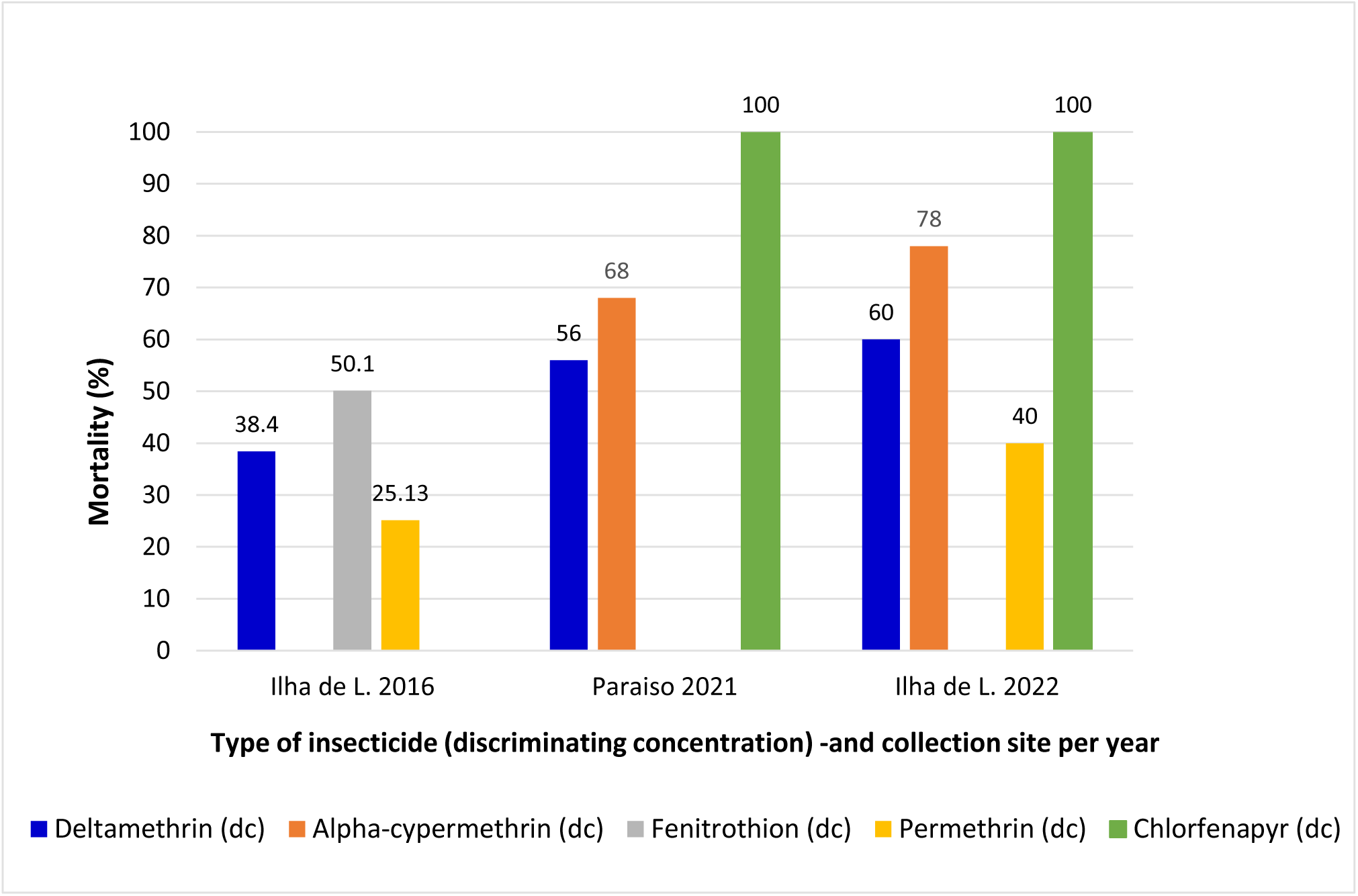
Susceptibility of *An. azevedoi* to different insecticides in sites of Luanda Province, Angola.

**Figure 3.**
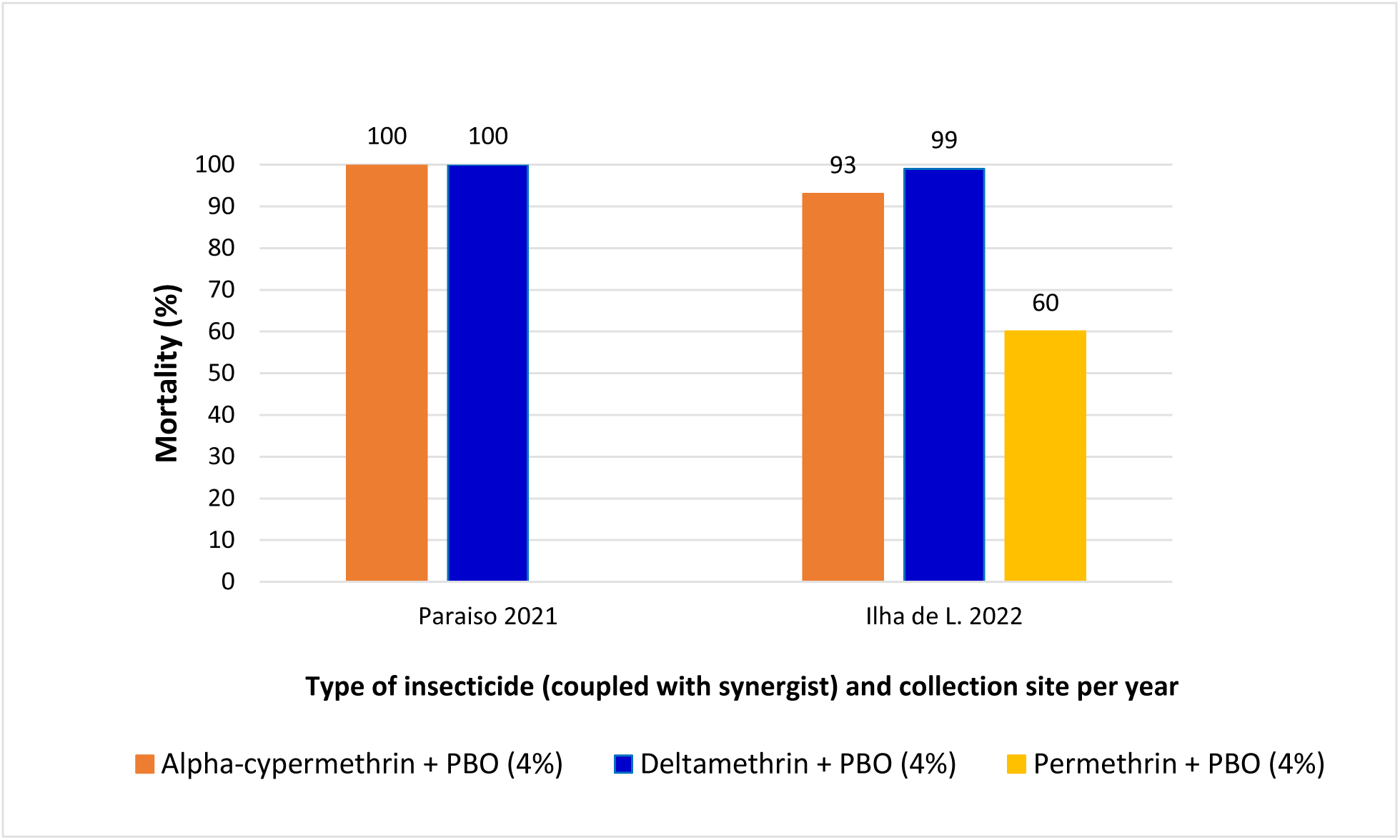
Susceptibility of *An. azevedoi* to pyrethroids with pre-exposure to a synergist (Piperonyl butoxide - PBO) in two sites of Luanda Province, Angola.

**Figure 4.**
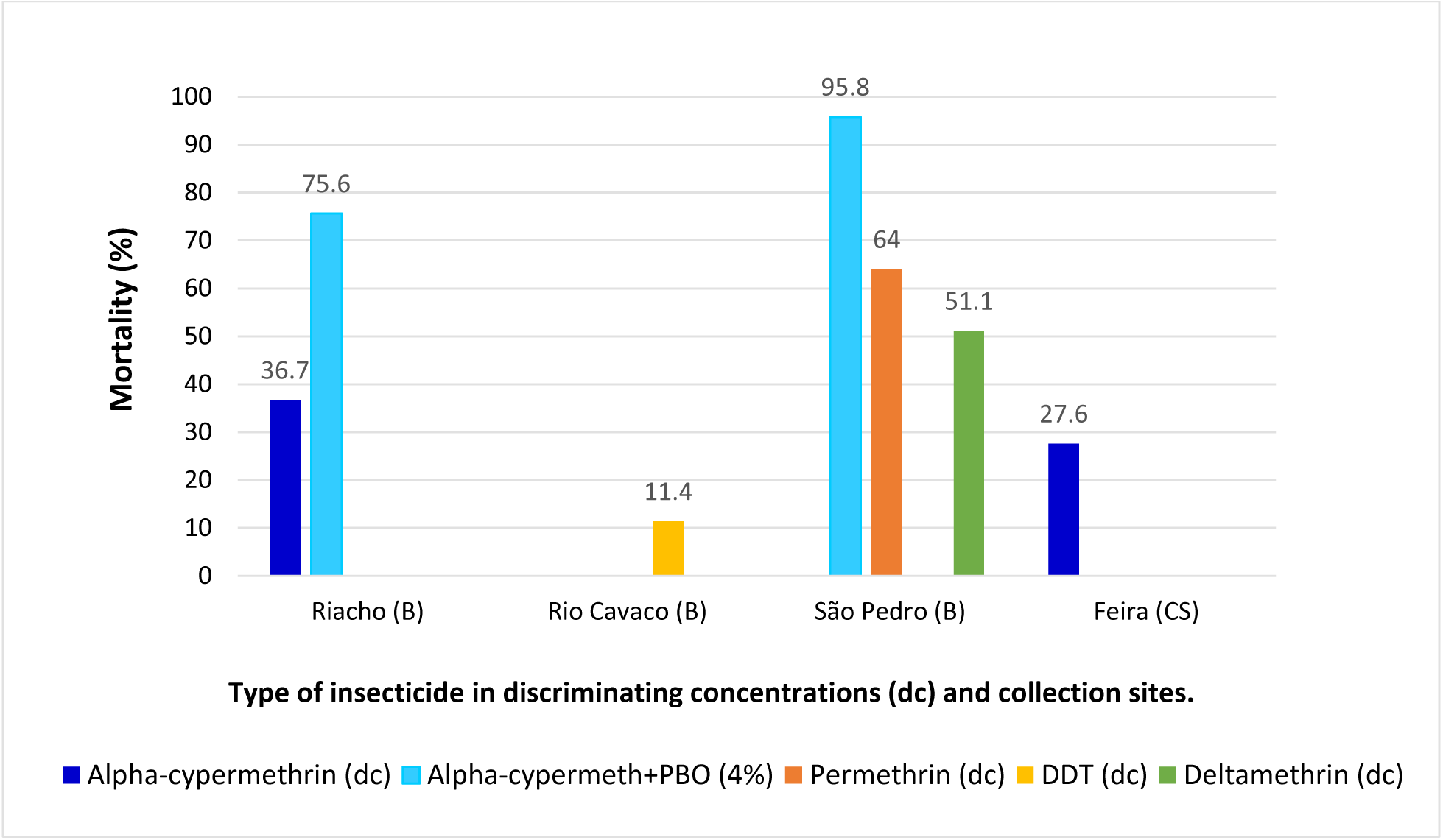
Susceptibility of *An. azevedoi* to public health insecticides in Benguela and Cuanza Sul provinces of Angola.

The relatively new insecticide chlorfenapyr (diagnostic dose of 100 ug/bottle) proved to be effective against the two populations of *An. azevedoi*, from Paraiso and Ilha de Luanda (Luanda Province), with tested individuals showing full susceptibility to this active ingredient (Figure 2).

For *An. azevedoi* populations from Luanda province (Ingombotas district) the recorded mortality was only 60% in individuals exposed to the discriminating concentration of permethrin, despite exposure to the synergist PBO (4%) in bioassays completed in 2022 (Figure 3).

Populations from Benguela and Cuanza Sul showed a generalized resistance to insecticides such as deltamethrin, permethrin, alpha-cypermethrin and DDT (Figure 4) in the corresponding sites.

In Cuanza Sul, the sampled populations were tested for only one insecticide, alpha-cypermethrin, at two dosages respectively, 0.05% (diagnostic dose) and 0.25% (5x) (35–36). Overall, the mosquito populations from Benguela and Cuanza Sul have confirmed insecticide resistance to three pyrethroids and to DDT, with only one population from Catumbela with “partially restored susceptibility” (95.8% mortality) to alpha-cypermethrin + PBO as per 2022 findings (Figure 4). Most of the outcomes representing the populations of *An. azevedoi* from these two western regions showed mortality levels under 80% (Figure 5) which are indicative of widespread resistance to pyrethroids and DDT.

**Figure 5.**
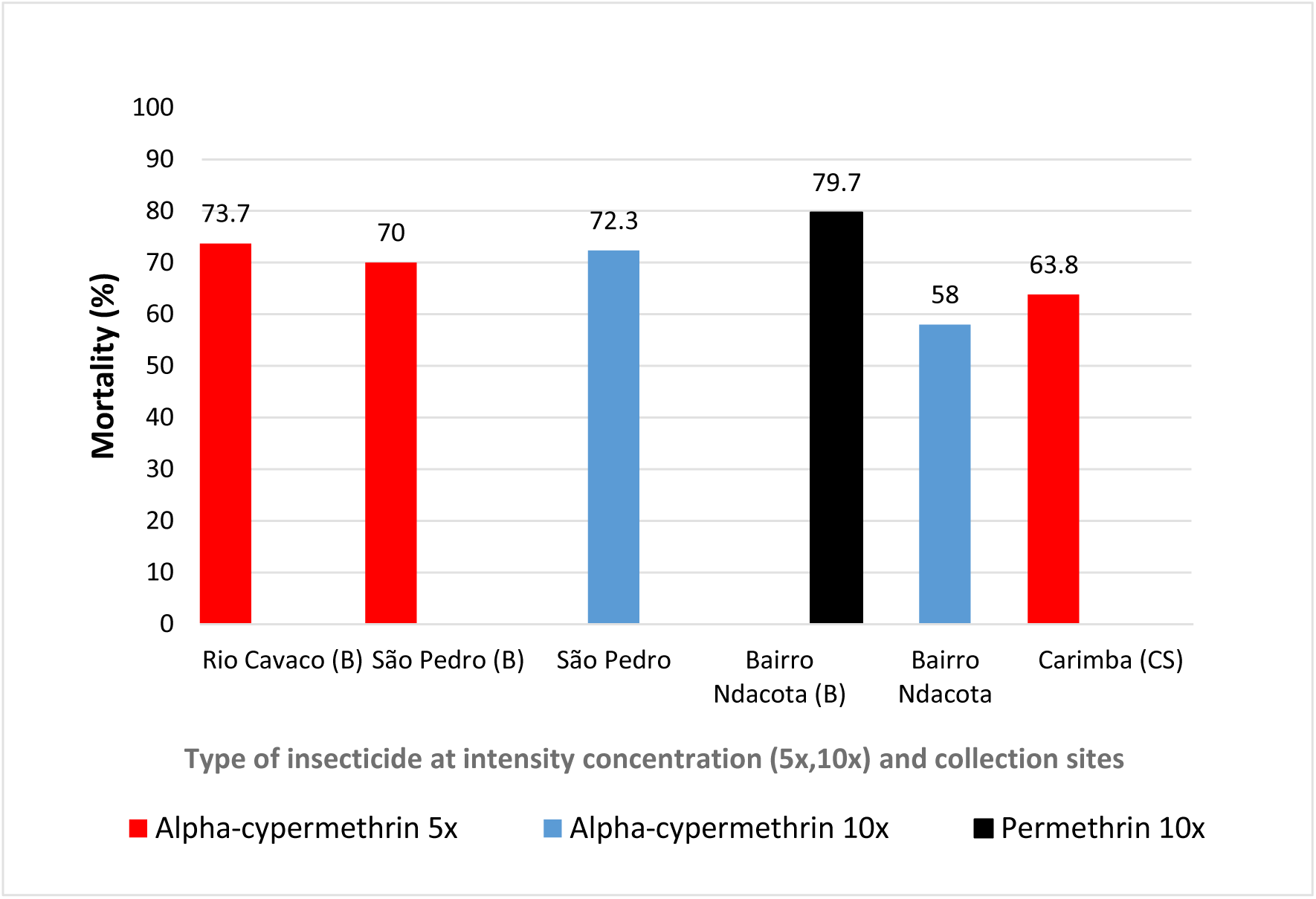
Susceptibility of *An. azevedoi* to pyrethroids using intensity assays (5x, 10x) in Benguela and Cuanza Sul provinces of Angola.

High resistance was documented in populations of *An. azevedoi* even when the bioassays included intensity doses equivalent to 5x and 10x the diagnostic concentration (Figure 5). For Rio Cavaco (Benguela), bioassays with DDT were also conducted with a smaller sample of mosquitoes (n=35), rendering the lowest mortality for the entire testing period (11.4%).

### Natural infectivity and blood source screening

The analysis of 26 *An. azevedoi* adult specimens captured in urban areas in Luanda, showed no evidence of *Plasmodium* infection, however, among these, nine specimens (seven of which were visible bloodmeals) were positive for the cytochrome b amplification indicating a group of mosquitoes positive for blood meals. Among these samples, three specimens were confirmed as fed on human blood with six specimen found inconclusive.

## DISCUSSION

Despite being a basic step in entomological surveillance, the identification of *Anopheles* species is a challenging task (24, 38) that relies on many variables, such as experience of the technician or professional conducting the taxonomic determination, availability of optical equipment and, the choice of an appropriate taxonomic key - in consideration of the geographical distribution of the *Anopheles* species being observed (39–41). It is worth noting that when following taxonomic keys (22, 24), the actual end point for the Angolan samples (later confirmed as *An. azevedoi*) is a duplet in which two species are highlighted: *An. azevedoi* and *An. listeri.* Thus, the essential question was to determine to which of these species did the Angolan samples belong to.

*Anopheles azevedoi* was first proposed as a new species occurring in Angola in 1969 by the Portuguese entomologist Henrique Ribeiro (15). Ribeiro highlighted the morphological similarity of *An. azevedoi* with *An. listeri* but also provided a thorough description of each stage while proposing additional characters to separate these two species when comparing adult females and males. This author also described internal morphology, specifically, male genitalia and female cibarium for both *An. azevedoi* and *An. listeri* (female cibarium referred as *pharynx*) (42).

At an earlier date, the morphological description of *An. listeri* was published by De Meillon (43), with the original description of the species. The holotype of *An. azevedoi* Ribeiro 1969, is associated to the coastal city of Angola named Moçâmedes (near the Atlantic Ocean), in the Province of Namibe, western region of the country; south of the provinces of Benguela, Cuanza Sul and Luanda. Comparatively, the type locality of *An. listeri* is the region of M’singa, Zululand in South Africa (43) which is in the eastern coast of the country, closer to the Indian Ocean coast (44).

Both species, *An. azevedoi* and *An. listeri*, are documented as tolerant to high salinity levels due to their occurrence along coastal environments (44). The distinction between the geographical range of these two species was also highlighted by Coetzee (24). In the main taxonomic key for the region (24) Coetzee described *An. azevedoi* as a species distributed in “south-western Africa only”, indicating the importance of the geographical origin of the specimens; and, for *An. listeri*, Coetzee included “southern Africa only” (24). Regional records of the geographical range of these two species show that *An. listeri* is widely distributed in Zimbabwe, South Africa, Namibia, Botswana, Mozambique and Southern Angola (21, 44), and *An. azevedoi*, has been documented in Angola, Namibia and South Africa (45). Though there is a potential overlapping in the geographical distribution of *An. listeri* and *An. azevedoi*, the existing literature indicates that along the Atlantic coast of Angola, the occurrence of *An. azevedoi* is clearly well documented and expected in the provinces sampled by our research teams (Benguela, Namibe, Luanda and Cuanza Sul). Our study confirms the geographical range previously highlighted by Ribeiro (16) and contributes with three new sites; two in the province of Luanda (Cacuaco and Ingombotas) and one in Benguela (Catumbela) (Figure 1).

In our current study, samples of *An. azevedoi* were not infected with *Plasmodium* sporozoites from specimens originally collected in Luanda province. However, wild specimens were found with human blood, which indicates that the species is anthropophilic. During field work, our teams linked breeding sites of the species within urban areas or in close proximity to them, specifically residential neighborhoods and beaches where the movement of people was recorded as constant (Luanda Province). The role of this species as a potential malaria vector remains unclear and will require further sampling to determine if populations of *An. azevedoi* have any role in malaria transmission. For now, our data and Ribeiro’s notes on the bioecology of the species (16), point to the fact that individuals of *An. azevedoi* are readily abundant in urban and rural areas of Angola, and feed on humans.

In Ribeiro’s work on the ecology of *An. azevedoi* (16) a wide geographical range is informed for the species, including Cuanza Sul (municipality of Porto Redondo, known today as Sumbe), Benguela (Benguela) and Namibe (municipalities of Porto Alexandre, São Nicolau, Mucuio and Saco do Girau). Ribeiro reported the species occurring only within 60 km of the Atlantic coast of western Angola. His records described high number of *An. azevedoi* individuals actively biting inside human habitation at late afternoon and dusk, exiting houses in the morning between 6:30 - 7:30 am. He documented *An. azevedoi* in resting locations with most females in the outdoors close to rocks and rock holes. Most of *An. azevedoi* breeding sites were associated with human activities, had high salt content, and abundance of filamentous algae; ranging from shaded to unshaded shallow pools. No infection with *Plasmodium* was documented in his surveys though dissections of salivary glands were performed (16).

The molecular confirmation of *An. azevedoi* was achieved by amplifying two genetic markers, one from the mitochondrial DNA, cytochrome oxidase (*COI*) gene, and a section of the ribosomal DNA, the internal transcribed spacer 2 (ITS2). Both *COI* and ITS2 are widely used in molecular studies and considered as reliable taxonomic markers to confirm *Anopheles* species (46–49). The availability of these markers is essential when only mosquito parts are recovered in the collections, and critical features are not available for unequivocally morphological identification. An added advantage of obtaining molecular data is the extensive database of reference sequences available for species identification of multiple taxa (46–49).

The closest species to *An. azevedoi*, when considering the morphological characters employed by classical taxonomy (24), is *An. listeri*. The latter has been documented with overlapping distribution with *An. azevedoi*, given that both species occur in the southern region of Africa. Despite records in earlier publications (16, 23) documenting immatures of both species in Angola, the accounts of *An. listeri* presence in the scientific literature is notably scarce, with the latest mentions of the species found in 2018 (50) with sites from South Africa, and an additional study in 2015 (51) with samples from Angola. Unfortunately, no DNA sequences were made available by either group of authors from these works.

Our search for available sequences of *An. listeri* rendered one entry in GenBank (accession number PQ154287). The ITS2 sequence (611 bp) corresponds to specimens of *An. listeri* from Angola and characterized in the supplementary data of said publication (one individual collected in 2010 with unspecific coordinates) (52). When comparing the two sequences, our Angolan sample (*An. azevedoi*) vs. PQ154287 specimen (*An. listeri*) (52), the two ITS2 sequences present 97% similarity, indicating clear species genetic differentiation. The general assumption by scholars around numerical BLAST outputs state that values for sequence similarity such as 98% or higher would be indicative of individuals belonging to the same species (53–55).

The only available *COI* sequences of *An. azevedoi* on GenBank are our study’s and that of a recent study also conducted in Angola (14). A mosquito survey in Angola (14) originally identified a specimen as part of the *Anopheles funestus* group by morphological characters but later corrected it by using *COI* (sequence made available in January 2026, under accession number PX837029). When specimen PX837029 (collected in Benguela) (14) was compared to the *COI* sequence generated by this study, the output showed 100% coverage and 99.85% similarity between them.

The outcomes from the insecticide surveillance are key to assessing the status and susceptibility of local mosquito populations to the common insecticides employed in public health operations, and especially relevant to decide which active ingredients should be considered for future distribution of insecticide-treated bed nets (ITNs) as part of a malaria control program. As per the case of Angola, the most recent ITN distribution campaign took place in 2022-2023, under projects named Health for All (HFA) and Procurement and Supply Management (PSM) led by MoHA. This ITN distribution campaign covered eight provinces of the country with the support from USAID/PMI and Global Fund. The campaign distributed single pyrethroid ITNs containing deltamethrin (commercial name is Yahe LN®) for provinces such as Cuanza Norte, Lunda Sul, Lunda Norte, Malanje, Uige and Zaire; these provinces are currently considered as hyperendemic for malaria transmission. In other regions of Angola, like Benguela and Cuanza Sul, the ITN campaign distributed two types of bed nets with single pyrethroid nets containing deltamethrin (commercial name PandaNet), alpha-cypermethrin (commercial name DuraNet), permethrin (commercial name Olyset), and deltamethrin with added piperonyl butoxide (PBO, a synergist; commercial name PermaNet® 3.0).

The outcomes of the insecticide susceptibility of local populations of *An. azevedoi* showed a generalized resistance to all the three pyrethroids tested in the sample sites of this study. Moreover, our findings suggest the presence of cross-resistance in tested populations of *An. azevedoi* between the organochloride (DDT) and all pyrethroids included in our study. This is explained by the mode of action of both classes of insecticides that act on the same target-site in the nerve membrane of the exposed insects, specifically the voltage-gated sodium channel interfering with the ion balance inside the insects’ cells (56).

Our data showed that only in three municipalities in which a synergist was employed, the susceptibility to deltamethrin and alpha-cypermethrin was recovered with 100% mortality of the mosquitoes exposed to both the synergist and insecticide (Figures 3-6). The synergist’s mode of action is the inhibition of enzymes associated with the metabolic detoxification of insecticides, recognized under the category of cytochrome P450 monooxygenases (57).

The generalized profile of high resistance to pyrethroids was also corroborated by HFA project when testing other species occurring in Angola, such as *Anopheles gambiae* s.s., and *Anopheles coluzzi*, with a set of common insecticides finding fixed resistance to all pyrethroids across six provinces across the Angolan territory (unpublished observations). HFA noted only a few exceptions where the exposure to PBO restored the susceptibility of *Anopheles* spp. populations to common pyrethroids. Among HFA’s final recommendations (unpublished observations) is the inclusion of dual active ingredient ITNs for future distribution campaigns in Angola, as part of core vector control strategies against malaria transmission.

A relevant outcome of the insecticide testing in our study was the susceptibility of populations of *An. azevedoi* to chlorfenapyr, a newly introduced insecticide. This insecticide, classified as a pyrrole, acts as a disruption of the ATP production thus causing loss of energy, cell dysfunction and death of target organisms (58).

Altogether, the results in this study inform that *An. azevedoi* is widely abundant but easily misidentified when only morphological characters are employed. Our findings link this species with urban environments such as residential areas and beaches. Though our study did not find specimens infected with *Plasmodium* sporozoites, three individuals sampled in a neighborhood of Luanda were documented with human blood. Considering the updated distribution and ecological considerations presented in our narrative, attention should be paid in future entomological surveillance programs so that new evidence is gathered to inform further presence and role of *An. azevedoi* in the public health scenario of Angola.

## ACKNOWLEDGEMENTS AND FUNDING

The authors appreciate the support and collaboration provided by the National Malaria Control Program (NMCP/ANGOLA) to conduct sampling of *Anopheles* specimens in different provinces of Angola. The authors are appreciative for Dr. John Gimnig’s support with the taxonomic determination of the adult samples received at the CDC - Atlanta. Dr. Adeline Chan, CDC Atlanta, previous PMI Entomologist in Angola coordinated entomological surveillance during 2015-2016 and secured the preservation of historical specimens from Angola. The funding for all technical activities came from various international donors such as the United States Agency for International Development (USAID) through the President’s Malaria Initiative that funded bilateral projects in Angola through Population Services International (PSI) acting as the implementing partner of USAID. PMI funded different projects with independent timeframes and workplans: AIRS (Africa Indoor Residual Spraying Project), VectorLink, and Health for All (HFA). The research activities conducted in Benguela and Cuanza Sul by the National Malaria Control Program, the Provincial Health Authorities, and The Mentor Initiative were funded by The Global Fund through the NMF3 Angola grant. Gonçalo Alves was funded in part by the Portuguese Foundation for Science and Technology PhD grant (2025.01126.BD). The funding agency had no role in the design of the study, the collection, management, analysis, and interpretation of the entomological data; the preparation, review, or approval of the manuscript; or the decision to submit it for publication. The study in Benguela and Cuanza Sul was approved by the Instituto Nacional de Investigação em Saúde de Angola (INIS). Letter reference: no. 12C.E/MINSA.INIS/2022. The findings and conclusions expressed herein are those of the author(s) and do not necessarily represent the official position of the US Agency for International Development (USAID), the U.S. President’s Malaria Initiative (PMI), or the Centers for Disease Control and Prevention (CDC).

## AUTHORS’ CONTRIBUTIONS

**GA, CM, VC, ADT, MSC, FE, CS, JP** conducted field collections, mosquito rearing, morphological determination and insecticide resistance bioassays. **TN, DC, JdR, AF, CP, JFM, MY**, **PM** and **CTG** contributed with study design and oversight of activities. **PM** conducted molecular testing, **AS** conducted ELISA testing and **GA** generated maps. **GA, CM, PM,** and **CTG** wrote initial drafts. **PM, GA, JP, MY** reviewed and contributed with feedback. **NMCP, PMI** authors supported project development and oversight of personnel and activities. **CTG, PM** and **GA** contributed to the final version of the manuscript.

## REFERENCES

1. The World Bank Group. 2022, Angola. Country climate and development report. 87pp.

2. Instituto Nacional de Estatistica (INE), Ministerio da Saude (MINSA) and ICF International (ICF). 2024. Inquérito de Indicadores Múltiplos e de Saúde de Angola, 2023–2024: Relatório de Indicadores Básicos. Luanda, Angola, se a publicação for feita em Luanda, Angola e Rockville, Maryland, EUA. https://dhsprogram.com/pubs/pdf/PR162/PR162.pdf [retrieved on August, 2025).

3. Shibre G. Social inequality in infant mortality in Angola: Evidence from a population based study. PLoS One. 2020;15(10):e0241049.

4. Cosep Consultoria, Consaúde, and ICF International. 2011. Angola Malaria Indicator Survey (MIS) 2011. Calverton, Maryland: Cosep Consultoria, Consaúde, and ICF International.

5. Haddad EA, Perobelli F, I. F-A. Uneven Integration: The Case of Angola. Preprint Research Square. 2020;10.21203/rs.3.rs-29035/v1.

6. World Health Organization (WHO). 2023. World Malaria Report 2023. https://www.who.int/teams/global-malaria-programme

7. Loughlin SO. The expanding Anopheles gambiae species complex. Pathogens and Global Health. 2020;114(1):1.

8. Carrara, G; Santolamazza, Federica; Fanello, C; Della Torre, A.; Petrarca, V. 2002. *-* In: Parassitologia. - ISSN 0048-2951. - STAMPA. - 44 (Suppl):(2002), pp. 43. (Presented at XXII Congresso Nazionale, Societa’ Italiana di Parassitologia, Grugliasco (Torino), June 11-14, 2002).

9. Cuamba N, Choi KS, Townson H. Malaria vectors in Angola: distribution of species and molecular forms of the Anopheles gambiae complex, their pyrethroid insecticide knockdown resistance (kdr) status and Plasmodium falciparum sporozoite rates. Malar J. 2006;5:2.

10. Calzetta M, Santolamazza F, Carrara GC, Cani PJ, Fortes F, Di Deco MA, et al. Distribution and chromosomal characterization of the Anopheles gambiae complex in Angola. Am J Trop Med Hyg. 2008;78(1):169–75.

11. Santolamazza F, Mancini E, Simard F, Qi Y, Tu Z, Della Torre A. Insertion polymorphisms of SINE200 retrotransposons within speciation islands of *Anopheles gambiae* molecular forms. Malar J. 2008;7(1):163.

12. Toto JC, Besnard P, Le Mire J, Almeida DS, Dos Santos MA, Fortes F, et al. [Preliminary evaluation of the insecticide susceptibility in Anopheles gambiae and Culex quinquefasciatus from Lobito (Angola), using WHO standard assay]. Bull Soc Pathol Exot. 2011;104(4):307–12.

13. Ribeiro H. Research on the mosquitoes of Angola. IV. Description of Anopheles (Cellia) azevedoi sp. nov. (Diptera, Culicidae). An Esc Nacl Saude Publica Med Trop (Lisb). 1969;3(1):113–23.

14. Alves G, Troco AD, Seixas G, Pabst R, Francisco A, Pedro C, et al. Molecular and entomological surveillance of malaria vectors in urban and rural communities of Benguela Province, Angola. Parasit Vectors. 2024;17(1):112.

15. Ribeiro H. Research on the mosquitoes of Angola. IV. Description of Anopheles (Cellia) azevedoi sp. nov. (Diptera, Culicidae). An Esc Nacl Saude Publica Med Trop (Lisb). 1969;3(1):113–23.

16. Ribeiro H. Research on the mosquitoes of angola africa part 5 on the bio ecology of anopheles azevedoi diptera culicidae. Garcia De Orta Serie de Zoologia. 1974;3(2):31–8.

17. Norris LC, Norris DE. Phylogeny of anopheline (Diptera: Culicidae) species in southern Africa, based on nuclear and mitochondrial genes. J Vector Ecol. 2015;40(1):16–27.

18. Correia CDN, Amraoui M, Santos JA. Assessment of Climate Change in Angola and Potential Impacts on Agriculture. Preprints. 2024.

19. Mesfin I, Benjamim M-H, Lebatard A-E, Saos T, Pleurdeau D, Matos J, et al. Evidence for Earlier Stone Age ‘coastal use’: The site of Dungo IV, Benguela Province, Angola. PLOS ONE. 2023;18(2):e0278775.

20. UNESCO. World Heritage Convention. Namibia National Commission for UNESCO, Reference 6094. Date of Submission 18/03/2016. Website https://whc.unesco.org/en/tentativelists/6094/ [retrieved December 2024 – title of page: Benguela current Marine Ecosystem Sites].

21. Gillies MT, De Meillon B. The Anophelinae of Africa south of the Sahara. Publications of the South African Institute for Medical Research, Johannesburg. 1968;54.

22. Gillies MT, Coetzee M. A supplement to the Anophelinae of Africa South of the Sahara. Publications of the South African Institute for Medical Research, Johannesburg. 1987;55:1–143.

23. Ribeiro H, Ramos HD. Guia ilustrado para a identificação dos mosquitos de Angola (Diptera. Culicidae). 1st ed. Lisboa: Boletim da Sociedade Portuguesa de Entomologia; 1995.

24. Coetzee MA. Key to the females of Afrotropical Anopheles mosquitoes (Diptera: Culicidae). Malaria Journal. 2020;19(1):70.

25. Dahan-Moss Y, Hendershot A, Dhoogra M, Julius H, Zawada J, Kaiser M, Lobo NF, Brooke BD, Koekemoer LL. Member species of the Anopheles gambiae complex can be misidentified as Anopheles leesoni. Malar Journal. 2020;19(1):89.

26. Koekemoer, L.L., Kamau, L., Hunt, R.H., Coetzee, M., 2002. A cocktail polymerase chain reaction assay to identify members of the Anopheles funestus (Diptera: Culicidae) group. Am J Trop Med Hyg 66, 804–811.

27. Wilkins, E.E., Marcet, P.L., Sutcliffe, A.C., Howell, P.I., 2009. Authentication scheme for routine verification of genetically similar laboratory colonies: a trial with *Anopheles gambiae*. BMC Biotechnol 9, 91.

28. Folmer O, Black M, Hoeh W, Lutz R, Vrijenhoek R. DNA primers for amplification of mitochondrial cytochrome c oxidase subunit I from diverse metazoan invertebrates. Mol Mar Biol Biotechnol. 1994;3(5):294–9.

29. Beebe NW, Saul A. Discrimination of all members of the Anopheles punctulatus complex by polymerase chain reaction-restriction fragment length polymorphism analysis. Am J Trop Med Hyg. 1995;53(5):478–81.

30. Camacho C, Coulouris G, Avagyan V, Ma N, Papadopoulos J, Bealer K, and Madden TL. 2009. BLAST+: architecture and applications. BMC Bioinformatics, 10, 421.

31. Malaria Research and Reference Reagent Resource Center (MR4). Methods in Anopheles Research. 2015 edition. Plasmodium circumsporozoite ELISA directions. Chapter 8. https://www.beiresources.org/Portals/2/VectorResources/2016%20Methods%20in%20Anopheles%20Research%20full%20manual.pdf (Accessed November 2025).

32. Sutcliffe AC, Irish SR, Rogier E, Finney M, Zohdy S, Dotson EM. Adaptation of ELISA detection of Plasmodium falciparum and Plasmodium vivax circumsporozoite proteins in mosquitoes to a multiplex bead-based immunoassay. Malar J. 2021;20(1):377.

33. Kent RJ, Norris DE. Identification of mammalian blood meals in mosquitoes by a multiplexed polymerase chain reaction targeting cytochrome B. Am J Trop Med Hyg. 2005;73(2):336–42.

34. World Health Organization (WHO). 2016. Test procedures for insecticide resistance monitoring in malaria vector mosquitoes, 2nd ed. World Health Organization. https://iris.who.int/handle/10665/250677

35. World Health Organization (WHO). 2022. Standard operating procedure for testing insecticide susceptibility of adult mosquitoes in WHO tube tests. Geneve; 2022.

36. World Health Organization (WHO). 2022. Standard operating procedure for determinining the ability of PBO to restore susceptibility of adult mosquitoes to pyrethroid insecticides in WHO tube tests. 01/14 January 2022. ISBN 978-92-4-004385-5.

37. World Health Organization (WHO). 2022. Standard operating procedure for testing insecticide susceptibility of adult mosquitoes in WHO bottle bioassays. Version: WHO Bottle-bioassay/01/14 January 2022. ISBN 978-92-4-004377-0.

38. Senjarini K, Abdullah MK, Azizah N, Septianasari MA, Tosin A, Oktarianti R, et al. Redesigning Primer of ITS2 (Internal Transcribed Spacer 2) for Specific Molecular Characterization of Malaria Vectors Anopheles Species. Med Arch. 2021;75(6):418–23.

39. Akeju AV, Olusi TA, Simon-Oke IA. Molecular identification and wing variations among malaria vectors in Akure North Local Government Area, Nigeria. Sci Rep. 2022;12(1):7674.

40. Birungi K, Mabuka DP, Balyesima V, Namukwaya A, Chemoges EW, Kiwuwa-Muyingo S, et al. Eave and swarm collections prove effective for biased captures of male Anopheles gambiae mosquitoes in Uganda. Parasit Vectors. 2021;14(1):281.

41. Kittichai V, Kaewthamasorn M, Samung Y, Jomtarak R, Naing KM, Tongloy T, et al. Automatic identification of medically important mosquitoes using embedded learning approach-based image-retrieval system. Sci Rep. 2023;13(1):10609.

42. Snodgrass RE. Principles of Insect Morphology: Cornell University Press; 1993.

43. De Meillon, B. Anopheles listeri s.n. Journal of the Medical Association of South Africa. 1931;5:482–483.

44. Abdulla-Khan R, Coetzee M, Hunt RH. Bionomics and cytogenetics of Anopheles seretsei Abdulla-Khan, Coetzee, and Hunt, a new species from northern Botswana. J Am Mosq Control Assoc. 1998;14(3):253–5.

45. Irish SR, Kyalo D, Snow RW, Coetzee M. Updated list of Anopheles species (Diptera: Culicidae) by country in the Afrotropical Region and associated islands. Zootaxa. 2020;4747(3):zootaxa.4747.3.1.

46. Collins FH, Paskewitz SM. A review of the use of ribosomal DNA (rDNA) to differentiate among cryptic Anopheles species. Insect Mol Biol. 1996;5(1):1–9.

47. Zomuanpuii R, Ringngheti L, Brindha S, Gurusubramanian G, Senthil Kumar N. ITS2 characterization and Anopheles species identification of the subgenus Cellia. Acta Trop. 2013;125(3):309–19.

48. Rathnayake RAS, Wedage WMM, Muthukumarana LS, De Silva B. Genetic diversity, phylogenetic and phylogeographic analysis of Anopheles culicifacies species complex using ITS2 and COI sequences. PLoS One. 2023;18(8):e0290178.

49. Calzolari M, Bellin N, Dottori M, Torri D, Di Luca M, Rossi V, et al. Integrated taxonomy to advance species delimitation of the Anopheles maculipennis complex. Sci Rep. 2024;14(1):30914.

50. Erlank E, Koekemoer LL, Coetzee M. The importance of morphological identification of African anopheline mosquitoes (Diptera: Culicidae) for malaria control programmes. Malar J. 2018;17(1):43.

51. Carnevale P, Toto JC, Besnard P, Santos MA, Fortes F, Allan R, et al. Spatio-temporal variations of Anopheles coluzzii and An. gambiae and their Plasmodium infectivity rates in Lobito, Angola. J Vector Ecol. 2015;40(1):172–9.

52. Boddé M, Makunin A, Teltscher F, Akorli J, Andoh NE, Bei A, et al. Improved species assignments across the entire Anopheles genus using targeted sequencing. Front Genet. 2024;15:1456644.

53. Lobo NF, St Laurent B, Sikaala CH, Hamainza B, Chanda J, Chinula D, et al. Unexpected diversity of Anopheles species in Eastern Zambia: implications for evaluating vector behavior and interventions using molecular tools. Sci Rep. 2015;5:17952.

54. St Laurent B, Cooke M, Krishnankutty SM, Asih P, Mueller JD, Kahindi S, et al. Molecular Characterization Reveals Diverse and Unknown Malaria Vectors in the Western Kenyan Highlands. Am J Trop Med Hyg. 2016;94(2):327–35.

55. Davidson JR, Wahid I, Sudirman R, Small ST, Hendershot AL, Baskin RN, et al. Molecular analysis reveals a high diversity of Anopheles species in Karama, West Sulawesi, Indonesia. Parasit Vectors. 2020;13(1):379.

56. Soderlund DM. Molecular mechanisms of pyrethroid insecticide neurotoxicity: recent advances. Arch Toxicol. 2012;86(2):165–81.

57. Syme T, Gbegbo M, Obuobi D, Fongnikin A, Agbevo A, Todjinou D, et al. Pyrethroid-piperonyl butoxide (PBO) nets reduce the efficacy of indoor residual spraying with pirimiphos-methyl against pyrethroid-resistant malaria vectors. Sci Rep. 2022;12(1):6857.

58. Raghavendra K, Barik TK, Sharma P, Bhatt RM, Srivastava HC, Sreehari U, et al. Chlorfenapyr: a new insecticide with novel mode of action can control pyrethroid resistant malaria vectors. Malar J. 2011;10:16.

